# Two-Dimensional NMR Lineshape Analysis

**DOI:** 10.1101/039735

**Authors:** Christopher Andrew Waudby, Andres Ramos, Lisa D. Cabrita, John Christodoulou

## Abstract

NMR titration experiments are a rich source of structural, mechanistic, thermodynamic and kinetic information on biomolecular interactions, which can be extracted through the quantitative analysis of resonance lineshapes. However, applications of such analyses are frequently limited by peak overlap inherent to complex biomolecular systems. Moreover, systematic errors may arise due to the analysis of two-dimensional data using theoretical frameworks developed for one-dimensional experiments. Here we introduce a more accurate and convenient method for the analysis of such data, based on the direct quantum mechanical simulation and fitting of entire two-dimensional experiments, which we implement in a new software tool, TITAN (TITration ANalysis). We expect the approach, which we demonstrate for a variety of protein-protein and protein-ligand interactions, to be particularly useful in providing information on multi-step or multi-component interactions.

## Introduction

Solution-state NMR spectroscopy is a powerful tool for the label-free characterization of structural and dynamical aspects of biomolecular interactions and equilibria^1,2^. Following changes in two-dimensional NMR spectra of macromolecules upon titration of a binding partner is a very common and information-rich approach capable of simultaneously characterizing thermodynamic (dissociation constant), kinetic (association and dissociation rates) and structural (chemical shift) aspects of interactions^3^. Critically, titration spectra are often sensitive probes of allosteric and multi-step binding mechanisms^4^, as used, for example, to elucidate the molecular mechanism underlying the remarkable selectivity of the chemotherapy drug Gleevec for inhibition of Abl tyrosine kinase^5^.

The appearance of NMR resonances during a titration experiment (e.g. to study a protein-ligand interaction) depends on the rate of exchange, *k*_ex_, between free and bound forms relative to the frequency difference, Δω, between these states^6^. When *k*_ex_≫Δω (‘fast exchange’), a progressive change in peak position is observed across the titration, while when *k*_ex_≪Δω (‘slow exchange’), separate free and bound resonances are observed with population-dependent intensities. Between these limiting cases (‘intermediate exchange’), more complex behaviour is observed in which chemical shift and intensity changes are not linearly related to the extent of binding. Analyses of chemical shift or intensity changes that neglect these effects can result in systematic errors in fitted *K*_d_ values^3^, but conversely, analyses that correctly account for the effects of exchange can extract valuable additional kinetic and mechanistic information on the system under investigation.

NMR lineshape analysis, also referred to as dynamic NMR, is a well-established method for the quantitative analysis of titration data based upon the fitting of one-dimensional spectra (or cross-sections from two-dimensional spectra) to theoretical or numerical solutions of the equations governing evolution of magnetization in an exchanging system^7-9^. As frequency differences, Δω, typically range from 10 to 10,000 s^−1^, NMR lineshape analysis can be suitable for the study of exchange processes, *k*_ex_, on timescales from 10 μs to 100 ms. The approach therefore strongly complements other NMR methods such as magnetisation exchange spectroscopy or relaxation dispersion^10-12^, as well as orthogonal biophysical techniques such as isothermal titration calorimetry^13,14^. Additionally, lineshape analysis can be a powerful probe of more complex reaction mechanisms^4^, such as cooperative or multi-step binding^15,16^, induced fit or conformational selection^17^, coupled folding and binding of intrinsically disordered proteins^11^, allostery^18,19^, enzyme catalytic cycles^9,20^ and ultrafast protein folding^21^. A variety of software packages have been described to implement the analysis^4,15,22^.

The extension of lineshape analysis to two-dimensional experiments, e.g. ^1^H,^15^N-HSQC or HMQC experiments, presents a number of additional features not encountered in one-dimensional experiments. Firstly, as distinct frequency (chemical shift) differences are associated with each dimension (Fig. 1a), the description of two-dimensional resonances as being in fast or slow exchange is not technically valid: lineshapes in each dimension may exhibit distinct behaviours (Fig. 1b-d). Secondly, relaxation occurring during the pulse sequence results in intensity changes that necessitates the normalisation of one-dimensional cross-sections^22^. As will be discussed below, this risks introducing both random and systematic errors into analyses. In addition, current analysis methods cannot be applied to experiments such as the HMQC, in which magnetisation is not single quantum during the indirect evolution period. Finally, we observe that the application of existing one-dimensional lineshape analysis methods has been severely limited by the problem of peak overlap, ubiquitous in spectra of complex biomolecules. In short, therefore, there is an urgent need for a theoretically rigorous (yet accessible) method for the analysis of two-dimensional datasets. In this manuscript, we describe such an approach, based on the direct simulation and fitting of two-dimensional spectra, which can fully account for the effects of exchange in common biomolecular NMR experiments, while efficiently handling the fitting of overlapping resonances. We anticipate that this approach will help facilitate more accurate and informative analyses of common titration experiments.

**Figure 1.**
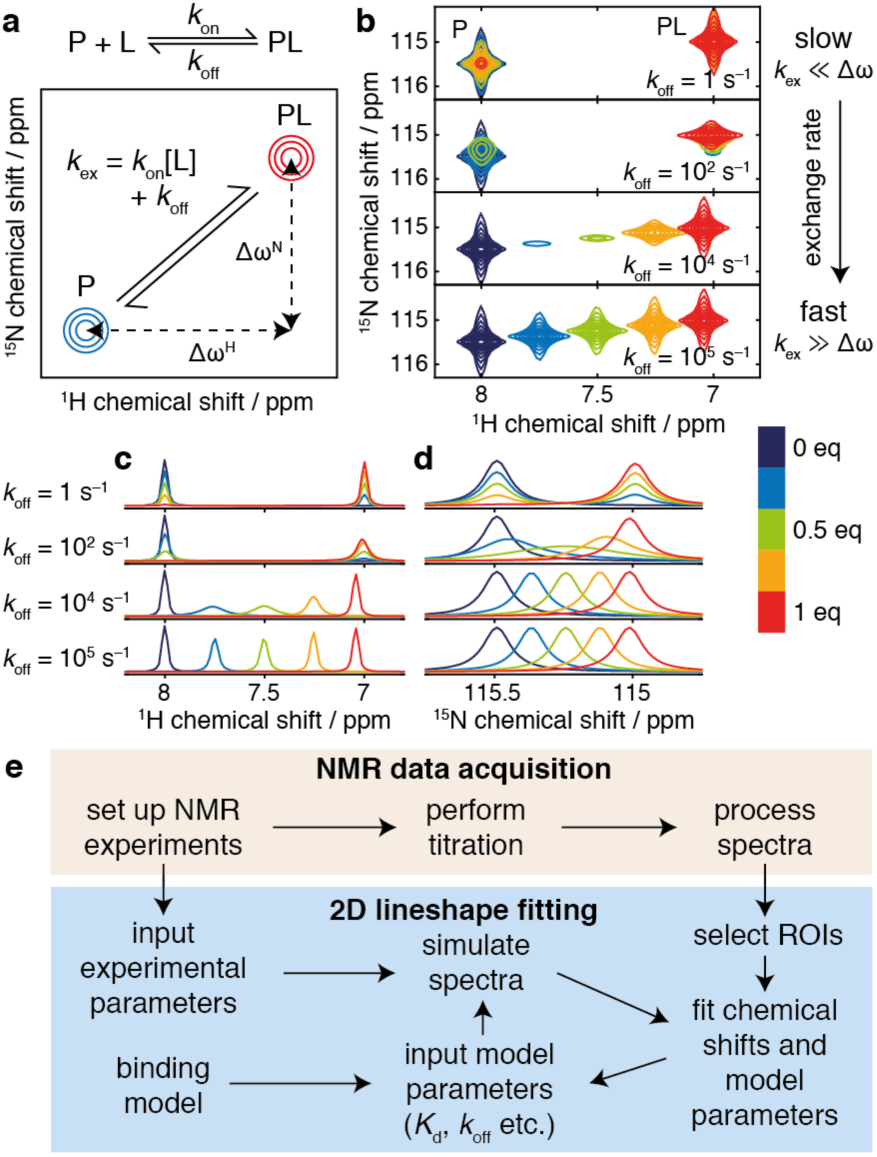
Principles of two-dimensional lineshape analysis. (a) Schematic showing the definition of the exchange rate and frequency differences for a two-state protein-ligand interaction. (b) Simulated ^1^H,^15^N-HSQC spectra for a protein-ligand interaction (700 MHz, 1 mM protein concentration, *K*_d_ 2 μM, Δω^H^ 4400 s^−1^, Δω^N^ 220 s^−1^) illustrating two-dimensional lineshapes that may arise under various exchange regimes. Contour levels are constant across all spectra. (c, d) ^1^H and ^15^N projections of HSQC spectra shown in Fig. 1b, normalised by integration. (e) Outline of the data acquisition process and the two-dimensional lineshape analysis procedure. ROIs, regions of interest.

## Results and Discussion

Existing approaches to lineshape analysis are based upon fitting solutions of the equations governing the evolution of magnetisation during chemical shift evolution periods to cross-sections of the observed spectra. In this manuscript, we propose extending this approach by calculating the evolution of magnetisation throughout the specific pulse sequence applied, by direct quantum mechanical simulation in Liouville space^23,24^ (see Methods). The two-dimensional interferograms thus obtained may be Fourier transformed to yield spectra suitable for comparison with experimental data. We have implemented this analysis using an optimised *in silico* ‘virtual spectrometer’ approach for the simulation and fitting of two-dimensional NMR experiments and datasets. This is configurable to match experimental acquisition parameters, allowing the efficient calculation of complete two-dimensional spectra against which best-fitting chemical shifts, linewidths and model parameters, such as binding constants and dissociation rates, can be determined using an iterative least-squares procedure (Figure 1e).

Our new approach brings several important advantages over one-dimensional methods, both in terms of convenience and accuracy. The direct analysis of two-dimensional spectra allows far greater flexibility in avoiding peak overlap, a problem ubiquitous in the congested spectra typical of biomolecules: for each spectrum, arbitrary regions of interest (ROIs) can be defined to exclude regions of peak overlap, or alternatively groups of overlapping resonances can be fitted simultaneously. In addition, the global fitting of multiple ROIs, all reporting on a common interaction as described below, provides a robust tool for monitoring the quality of fits. Also, by tracking the relaxation (decay) of magnetisation across the entire pulse sequence, calculations can fully account for the differential relaxation of states during execution of the pulse program. Such effects arise frequently in slow-intermediate exchange regimes when the various states (conformations) of the macromolecule do not have the same linewidth (for example in folding/unfolding reactions, or dimerization and other association/dissociation reactions), and can induce systematic errors when using one-dimensional analysis methods as the amount of magnetization associated with a particular state is no longer proportional to its population (Supplementary Fig. S1). A similar effect can also distort one-dimensional analyses of HMQC experiments, due to the influence of ^1^H chemical shift changes on multiple quantum coherences during the indirect detection period (Supplementary Fig. S2). Lastly, and again because relaxation is fully treated throughout the pulse sequence, the intensity of NMR signals can be rigorously compared between titration spectra. In contrast, one-dimensional methods require that every peak cross-section must be individually normalised (Fig. 1b-d), either by integration, which may introduce large errors due to noise in the spectrum, or by fitting, which introduces a large number of additional degrees of freedom, ultimately resulting in a less powerful analysis.

The two-dimensional lineshape analysis method described here has been implemented in the software package TITAN (TITration ANalysis, http://www.nmr-titan.com). TITAN can be used to simulate the HSQC and HMQC pulse programs commonly used to monitor protein-ligand interactions (Supplementary Fig. S3, Table S1), and data can be fitted to a range of binding models, from simple two-state interactions to more complex induced fit or conformational selection mechanisms. Example data and analysis scripts are provided, and a flexible ‘plug-in’ approach allows the implementation of additional pulse programs or binding models if required. Simple functions and interfaces are provided for the import of data, selection of ROIs, and global fitting and error analysis.

It is of paramount importance that parameters estimated by the lineshape analysis methods we describe are accompanied by reliable estimates of their experimental uncertainty. To this end, we have investigated the application of a bootstrap error analysis method based on resampling of fitting residuals in two-dimensional blocks(Kunsch, 1989) (Fig. 2a). In contrast to conventional methods based on resampling of individual points, this approach accounts for correlation between neighboring points, resulting in a more accurate estimation of parameter uncertainties. This method is also useful for linewidth measurements in single spectra. To validate the analysis, we generated test data for a two-state binding interaction in which the *K*_d_, *k*_off_ and noise level were systematically varied over several orders of magnitude (examples of which are shown in Fig. 2b). Parameter uncertainties were calculated by residual resampling using either conventional methods or 5×5 blocks. The distributions of the resulting *z*-scores (*z* = (*x*_fit_–*x*_true_) / σ_x_) are examined in Fig. 2c. While conventional residual resampling results in systematic underestimation of uncertainties, the distribution obtained by block resampling is close to a standard normal distribution, providing strong evidence that parameter values, and their associated uncertainties, are being correctly determined.

**Figure 2.**
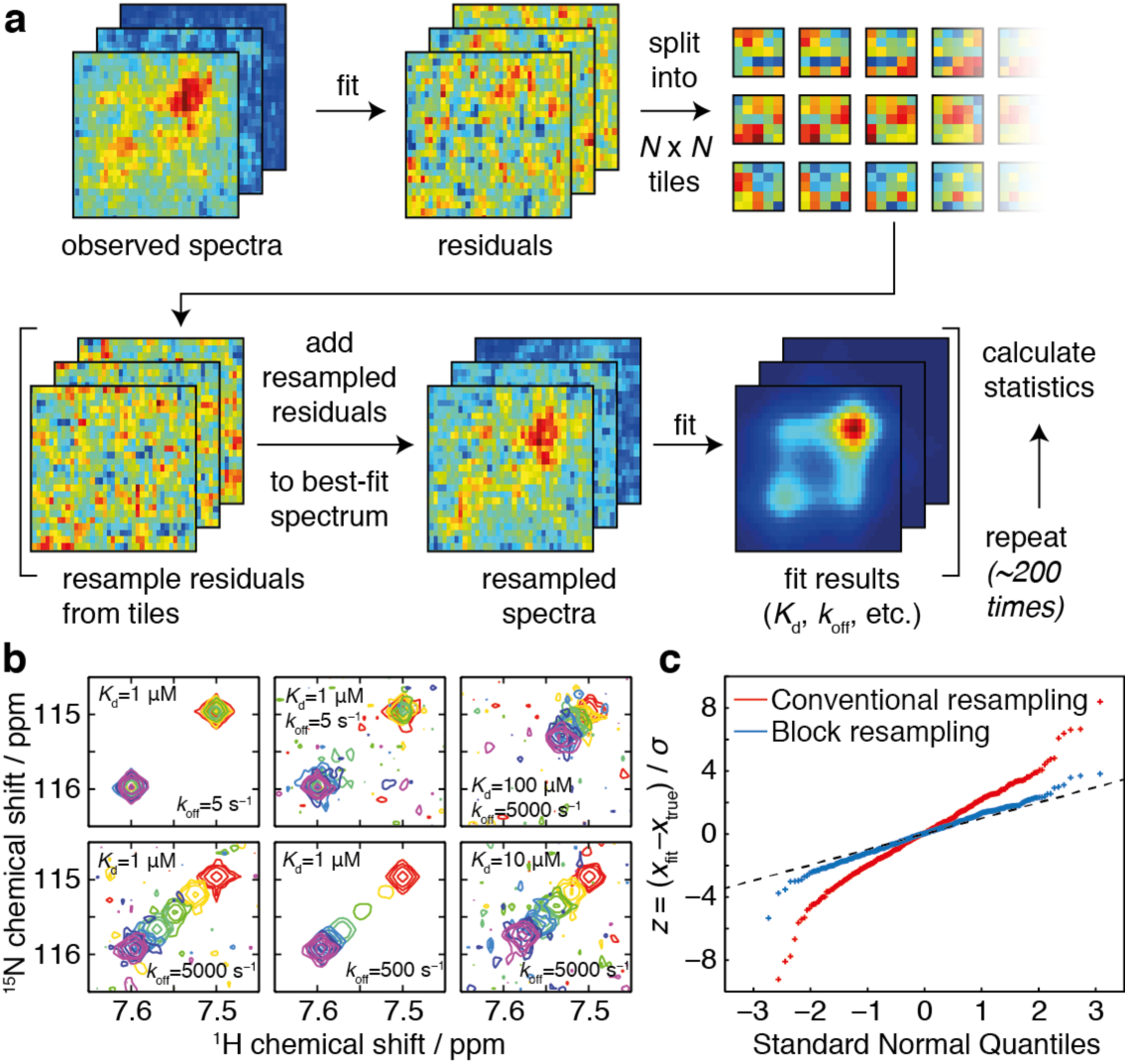
Error analysis and validation. (a) Implementation of error analysis using a block residual resampling scheme. (b) Simulated protein-ligand titrations with a fixed protein concentration of 50 μM and ligand concentrations of 0 (red), 12.5 (yellow), 25 (green), 50 (cyan), 62.5 (blue) and 75 μM (purple), with the *K*_d_ and *k*_off_ parameters varied as indicated. (c) Q-Q plot of *z*-scores of fitted parameters for the simulated test data in (b), with standard errors calculated by residual resampling using conventional methods or 5×5 blocks as indicated. The standard normal distribution is indicated by a dashed line.

We first applied TITAN to the analysis of previously reported NMR titration data: the interaction between the FIR RRM1-RRM2 protein and the FBP and FBP3 Nbox peptides, two key components of the FUSE system for regulation of c-myc transcription during the cell cycle^25^. For each titration series, ca. 30 FIR resonances were fitted globally to a two-state binding process (Fig. 3a-b, Supplementary Fig. S4–5). The fitted binding constants were consistent with those originally reported (Fig. 3c). Critically, the interaction kinetics were also determined, from which it may be observed that the stronger affinity of FBP Nbox is mainly due to the increased lifetime of the bound state (370 μs vs 67 μs), rather than to more rapid association. This finding highlights that disrupting the functional interaction is better achieved by reducing the lifetime of the complex rather than acting on the association of the two molecules, which may narrow the focus in the design of compounds to manipulate the FBP-FIR interaction. A further analysis of the functional interaction of FIR with oligonucleotides from the FUSE target DNA^25^ also shows the results of two-dimensional lineshape fitting to be in good agreement with previous determinations (Supplementary Fig. S6). Overall, these results both validate the analysis procedure and illustrate the general ability to analyse and extract new results (e.g. binding kinetics) from existing datasets.

**Figure 3.**
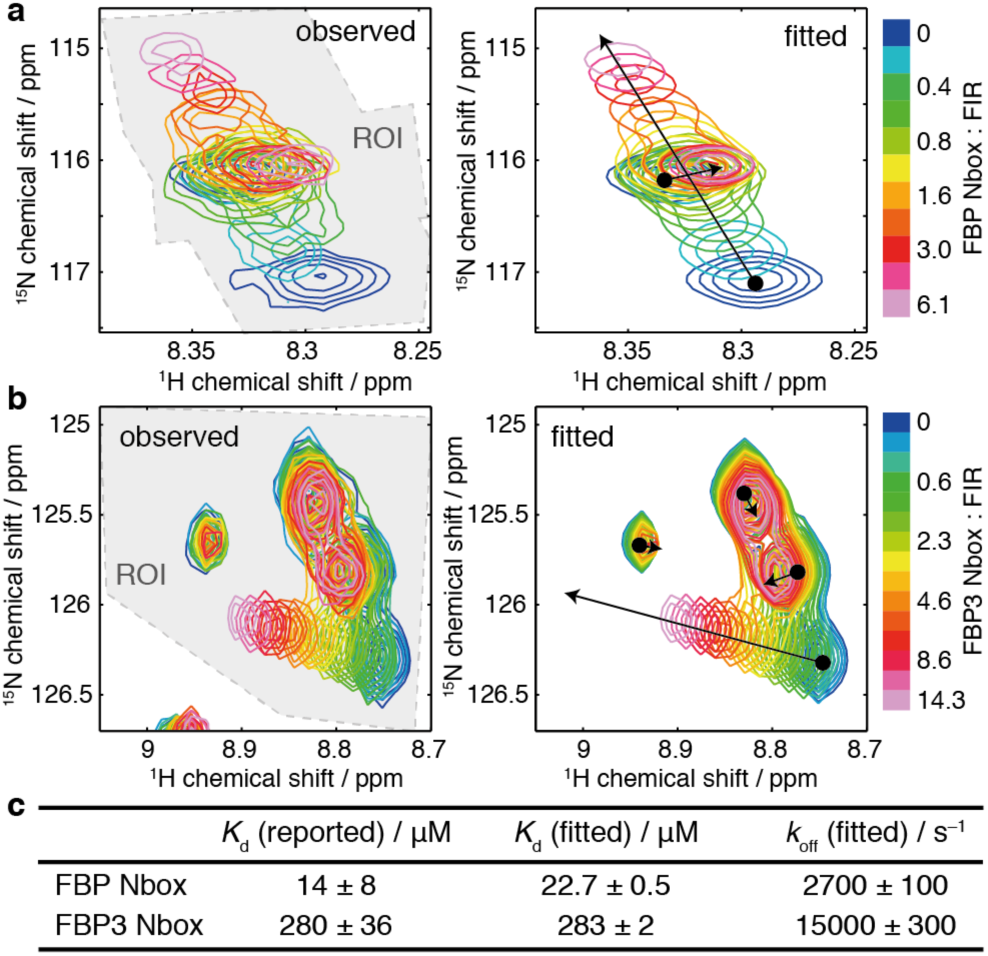
Analysis of the interaction of FIR RRM1-RRM2 with Nbox peptides. (a) Observed and fitted ^1^H,^15^N-HMQC spectra of 41 μM FIR RRM1-RRM2 upon titration of FBP Nbox. Shaded area indicates the selection region of interest (ROI). (b) Observed and fitted ^1^H,^15^NHMQC spectra of 41 μM FIR RRM1-RRM2 upon titration of FBP3 Nbox. (c) Reported and fitted binding model parameters for the interaction of FIR RRM1-RRM2 with FBP and FBP3 Nbox.

As discussed above, NMR lineshapes are a sensitive tool for identifying and investigating multi-step interaction mechanisms such as induced fit or conformational selection^4,5^ (Supplementary Fig. S7). However, the dependence of lineshapes on binding mechanisms can be non-intuitive, and so example TITAN scripts are provided that allow users to easily explore mechanisms and ranges of parameters of relevance to particular systems. Moreover, we have also developed an interactive online application that allows the rapid exploration of the most common binding models using a simple graphical user interface (accessible at http://www.nmr-titan.com).

As an example of the analysis of a more complex binding mechanism, we have investigated the interaction of calmodulin (CaM) with the drug trifluoperazine^26^ (TFP). Crystal structures have been determined with 1, 2 and 4 equivalents of TFP bound^27-29^, while previous NMR studies have observed complex patterns of chemical shift changes that have hitherto precluded quantitative analysis^26^. In particular, the direction of chemical shift changes were observed to change across the titration, which is indicative of a sequential binding mechanism (Fig. 4a-b, Supplementary Fig. S7). To obtain a quantitative model of this interaction, we recorded ^1^H,^15^N-HSQC spectra across a titration of uniformly ^15^N-labelled (Ca^2+^)_4_-CaM with TFP, and attempted to fit the data to a model describing the sequential binding of 4 TFP molecules (Fig. 4a, Supplementary Fig. S8). We found that this minimal model, based on the simultaneous fitting of 33 residues evenly distributed around the protein, described the observed data accurately, revealing a hierarchy of binding constants together with their associated rate constants (Fig. 4b). Moreover, the fitted chemical shift changes provide useful structural information on the various binding sites: when projected onto previously determined crystal structures we found that the pattern of chemical shift changes reproduced the crystallographic order of the multiple binding sites, and could be used to resolve the order of the third and fourth binding sites (Fig. 4c-f).

**Figure 4.**
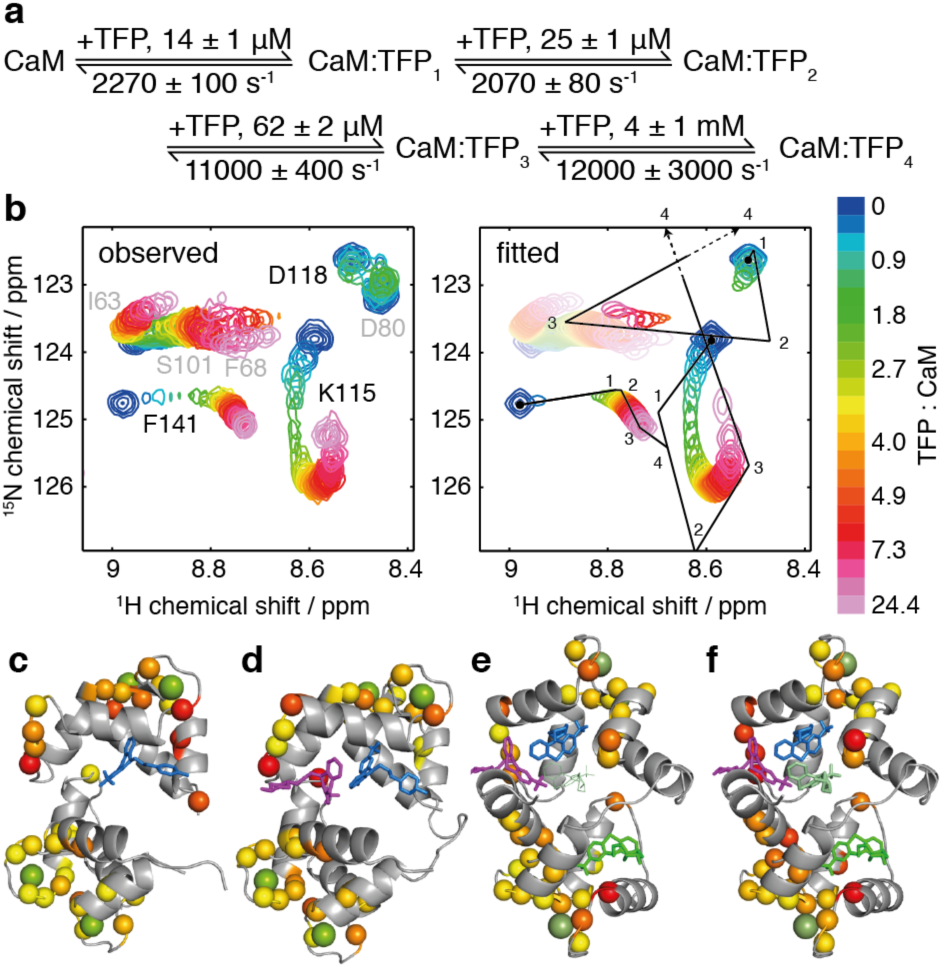
Analysis of the interaction of Ca^2+^_4_-CaM with TFP. (**a**) Sequential binding model showing fitted dissociation and rate constants. (**b**) Observed and fitted ^1^H,^15^N-HSQC spectra of 40 μM Ca^2+^_4_-CaM upon titration of TFP. Chemical shift changes are shown for K115, D118 and F141. D80 was excluded from the fitted region of interest, and contours of the fitted I63, F68 and S101 resonances are desaturated to improve contrast for the remaining resonances. (**c-f**) Chemical shift differences (Δδ=(Δδ_H_^2^+(Δδ_N_/5)^2^)^1/2^) between bound states, projected onto CaM-TFP crystal structures. Green spheres indicate calcium atoms. (**c**) 0–1 eq, pdb 1CTR^27^, 0–0.25 ppm yellow–red; (**d**) 1–2 eq, pdb 1A29^29^, 0–0.4 ppm yellow–red; (**e**) 2–3 eq, pdb 1LIN^28^, 0–0.7 ppm yellow–red; and (**f**) 3–4 eq, pdb 1LIN, 0–1 ppm yellow–red.

In summary, we have presented an improved method to extract structural, thermodynamic and kinetic information on protein-ligand interactions using two-dimensional NMR spectroscopy. As a proof of principle we have applied the method to two very different systems, the 1:1 interaction between the FBP and FIR c-*myc* transcriptional regulators, and the multi-state interactions between the drug TFP and calmodulin. In both cases, we show that our analysis yields novel structural and mechanistic insight into the interactions. The method is applicable to the analysis of a wide range of processes and systems. Direct quantum mechanical simulation of experiments provides a flexible approach that is extensible to more complex pulse sequences (for example the CPMG-HSQC experiment^30^, Supplementary Fig. S9). The analysis can also be applied to more complex spin systems. For example, provided that fast-relaxing coherences can be neglected^31^, methyl-TROSY ^1^H,^13^CHMQC measurements of CH_3_ groups can be treated as two-spin systems using the existing HMQC implementation. This will extend the use of TITAN to the study of high molecular weight systems. Ultimately, we expect these methods to facilitate the routine quantitative analysis of NMR titration data to resolve aspects of complex interaction mechanisms.

## Methods

### Code availability

The TITAN application and source code (developed and tested in MATLAB 2015b) is freely available for academic use from http://www.nmr-titan.com. An interactive online tool for the exploration of common binding models (developed in Mathematica 10.2, Wolfram Research Inc., Champaign, Illinois) is freely available at the same address.

### Two-dimensional lineshape analysis procedure

Fitting is performed as outlined in Fig. 1e. Firstly, a pulse program is specified together with associated spectral parameters, such as the number of scans, number of points, sweep widths, operating frequency and apodization. A series of titration points is set up, in which protein and ligand concentrations are specified, and a binding model selected (in order to convert concentrations and global parameters such as *K*_d_ and *k*_off_ into appropriate exchange superoperators). Next, sets of spin systems are created, specifying initial estimates of peak positions for each state, and for each spin system regions of interest (ROIs) are selected for each spectrum, using the graphical user interface provided. Only data within these ROIs are used for fitting, which provides a simple means to avoid regions of peak overlap, although in many cases it is also effective to use larger ROIs and fit the overlapping peaks directly. The fitting process itself is best conducted as an iterative process, due to the large number of free parameters (each spin has two chemical shifts and linewidths associated with each state, plus global model parameters). For example, it is often effective to fit chemical shifts and linewidths for the first spectrum alone, then hold these parameters constant for the remainder of the session. If additional constraints are known, for example *K*_d_ values from other biophysical methods, these can also be held constant. Finally, when a satisfactory fit is obtained (simple functions are provided for the two and three dimensional visualisation and inspection of fits), bootstrap error analysis can be performed, from which parameter uncertainties and covariances are determined.

### Simulation of two-dimensional spectra

Two-dimensional spectra are simulated by propagation of density operators in a composite Liouville space formed from the direct product of the chemical state space and the spin Liouville space^7,23,24^, incorporating an exchange super operator, *K*, derived from the specified binding model and calculated at each point in the titration series. For example, in the case of three-state exchange, the form of *K* is:

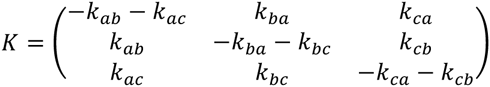

A vector of equilibrium concentrations, *p*_0_, is derived from the microscopic rate constants and this is used to form an initial density operator, *ρ*_0_ (Supplementary Table S1). This is then propagated for all required values of the indirect evolution period, *t*_1_, until the start of the direct acquisition period. To accelerate calculations only active subspaces are propagated, and the effect of some pulses is therefore to rotate between these subspaces (Supplementary Fig. S1). Basis sets and superoperators are tabulated in Supplementary Table S1. To implement frequency discrimination, cosine and sine modulated amplitudes are obtained simultaneously as real and imaginary components (Supplementary Fig. S1, Table S1), and observable magnetisation at the point of acquisition is then mapped onto chemical states by the operator *M*_+_ (Table S1). Thus, for each spin, σ, we obtain a complex-valued (*n*_1_ × *k*) matrix *A*_σ_, where *n*_1_ is the number of complex points and *k* is the number of states.

Next, for each spin lineshapes are calculated in the direct dimension via the McConnell equations^6,8^, e.g. in the case of three-state exchange:

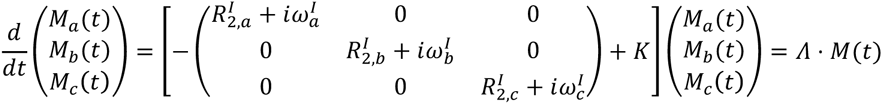

The evolution operator *Λ* is diagonalised, such that each eigenvalue λ*_*i*_* of *Λ* represents the complex frequency (i.e. chemical shift and linewidth) of a Lorentzian resonance. From the eigenvectors we can determine the amplitudes of each eigenstate associated with starting populations of magnetisation in pure chemical states, and thus we can compute a (*k* × *n*_2_) matrix, *B*_σ_, containing the combinations of Lorentzian lineshapes that originate from these pure states, where *n*_2_ is the number of (frequency domain) points in the direct dimension. We note that, if in the future the simulation of scalar coupled systems is required, the calculation at this stage could also be performed using a quantum mechanical density operator formalism as above.

The complete two-dimensional dataset can then be calculated, summing across all fitted spin systems, as *Y*(*t*_1_, *ω*_2_) = ∑_*σ*∈spins_*A*_*σ*_*B*_*σ*_. A window function can be applied directly to the indirect dimension, while in the direct dimension apodization and, if required, a uniform ^3^*J*_HNHA_ coupling, is introduced by convolution. Finally, the spectrum is Fourier transformed in the indirect dimension, with zero filling as required.

### One-dimensional lineshape fitting

^1^H and ^15^N lineshapes were obtained by integration over a rectangular region of interest, and these were fitted simultaneously to numerical solutions of the McConnell equations, with normalisation factors fitted for each spectrum, as previously described^20,22^.

### Validation by analysis of synthetic data

Protein-ligand titrations were simulated with a fixed protein concentration of 50 μM and ligand concentrations of 0, 12.5, 25, 50, 62.5 and 75 μM, with *K*_d_ values varied between 1 and 100 μM and *k*_off_ values between 5 and 5000 s^−1^. The performance of the two-dimensional fitting algorithm was investigated with different levels of noise in the synthetic dataset, and the uncertainties in the fitted values were determined by bootstrapping using standard residual resampling, and by resampling of 5×5 blocks.

### Analysis of FIR RRM1-RRM2 interactions with FBP Nbox, FBP3 Nbox and oligonucleotides

Titration data, as previously described^25^, were processed with exponential line broadening using nmrPipe^32^, then imported into MATLAB for analysis with TITAN. Data were fitted to a two-state ligand binding model in a two-stage process: chemical shifts and linewidths of the free state were determined using the first spectrum only, then chemical shifts of the bound state, linewidths of all states, and the binding model parameters *K*_d_ and *k*_off_ were fitted using the entire dataset. Error estimation was performed by residual resampling using 200 replicas and a 5×5 block size.

### CaM-TFP titration and analysis

Following previous protocols^26^, a 41 μM sample of uniformly ^15^N-labelled rat calmodulin was prepared in 10 mM imidazole (pH 6.5), 100 mM KCl, 100 μM EDTA, 5 mM CaCl_2_, 10% (v/v) D_2_O, 0.001% (w/v) DSS, and titrated with 5 mM or 50 mM stocks of TFP in an identical buffer to give TFP:CaM ratios of 0, 0.30, 0.61, 0.91, 1.22, 1.52, 1.83, 2.13, 2.44, 2.74, 3.05, 3.35, 3.66, 3.96, 4.27, 4.57, 4.88, 5.49, 6.10, 7.32, 9.76, 14.63 and 24.39. At each point, FHSQC experiments were acquired (298 K, spectral width 18 × 31 ppm, acquisition times 107 × 32 ms, 16 scans, 1.5 s recycle delay) using a Bruker Avance III NMR spectrometer operating at 800 MHz. Spectra were referenced to internal DSS^33^ and processed with 4 Hz and 10 Hz exponential line broadening in the direct and indirect dimensions respectively using nmrPipe^32^. Processed spectra were fitted to a five-state sequential binding model in stages:unbound chemical shifts and linewidths were determined using the first spectrum only, then all other chemical shifts and binding model parameters were fitted using the entire dataset. Given the conformations of the various protein-TFP complexes were not expected to vary significantly, to reduce the number of free parameters resonance linewidths were fitted as shared parameters, equal across all states of the model. Error estimation was performed by residual resampling using 200 replicas and a 5×5 block size, performed in parallel using the UCL Legion high performance computing facility.

## Acknowledgements

We acknowledge the use of the UCL Biological NMR Facility, the UCL Legion High Performance Computing Facility (Legion@UCL), and the MRC for access to the Biomedical NMR Centre at the Francis Crick Institute, London. The research was supported by a Welcome Trust Investigator Award 097806/Z/11/Z (C.A.W., J.C.).

### Author contributions

C.A.W. conceived the analysis strategy, wrote the code, designed and performed experiments, and analyzed data. A.R. provided experimental data and edited the manuscript. L.D.C. provided reagents. C.A.W. and J.C. wrote the manuscript.

### Competing financial interests

The author(s) declare no competing financial interests.

## SUPPLEMENTARY INFORMATION

Table S1 Superoperators for calculation of evolution during pulse sequences.

Figure S1 Comparison of 1D and 2D lineshape fitting showing effect of differential relaxation.

Figure S2 Comparison of exchange lineshapes in HSQC and HMQC experiments.

Figure S3 Analysis of pulse sequences for lineshape analysis.

Figure S4 Fit results for the titration of FIR RRM1-RRM2 with FBP Nbox.

Figure S5 Fit results for the titration of FIR RRM1-RRM2 with FBP3 Nbox.

Figure S6 Fit results for the titration of FIR RRM1-RRM2 with oligonucleotides.

Figure S7 Simulation of binding mechanisms.

Figure S8 Fit results for the titration of Ca^2+^-CaM with TFP.

Figure S9 Comparison of exchange lineshapes in HSQC and CPMG-HSQC experiments.

**Table S1.** Superoperators for calculation of evolution during HSQC, HMQC and CPMG-HSQC pulse sequences (as depicted schematically in Fig. S3). *K* is the exchange matrix, *k* is the number of states, and *I_n_* is the *n* × *n* identity matrix.

|  | HSQC | HMQC | CPMG-HSQC |
| --- | --- | --- | --- |
| $\rho_0$ | $p_0 \otimes (0 \ 1 \ 0 \ 0)'$ | $p_0 \otimes (0 \ 1 \ 0 \ 0)'$ | $p_0 \otimes (0 \ -1 \ 0 \ 0)'$ |
| basis A | $\{I_x, I_y, 2I_x S_z, 2I_y S_z\}$ | $\{I_x, I_y, 2I_x S_z, 2I_y S_z\}$ | $\{I_x, I_y, 2I_x S_z, 2I_y S_z\}$ |
| basis B | $\{S_x, S_y, 2I_z S_x, 2I_z S_y\}$ | $\{ZQ_x, ZQ_y, DQ_x, DQ_y\}$ | $\{S_x, S_y, 2I_z S_x, 2I_z S_y\}$ |
| $L_{free,A}$ | $\bigoplus_{i=1}^K \begin{pmatrix} -R_{2,i}^I & -\omega_i^I & 0 & -\pi J \\ \omega_i^I & -R_{2,i}^I & \pi J & 0 \\ 0 & -\pi J & -R_{2,i}^I & -\omega_i^I \\ \pi J & 0 & \omega_i^I & -R_{2,i}^I \end{pmatrix} + K \otimes I_4$ | $\bigoplus_{i=1}^K \begin{pmatrix} -R_{2,i}^I & -\omega_i^I & 0 & -\pi J \\ \omega_i^I & -R_{2,i}^I & \pi J & 0 \\ 0 & -\pi J & -R_{2,i}^I & -\omega_i^I \\ \pi J & 0 & \omega_i^I & -R_{2,i}^I \end{pmatrix} + K \otimes I_4$ | $\bigoplus_{i=1}^K \begin{pmatrix} -R_{2,i}^I & -\omega_i^I & 0 & -\pi J \\ \omega_i^I & -R_{2,i}^I & \pi J & 0 \\ 0 & -\pi J & -R_{2,i}^I & -\omega_i^I \\ \pi J & 0 & \omega_i^I & -R_{2,i}^I \end{pmatrix} + K \otimes I_4$ |
| $L_{free,B}$ | $\bigoplus_{i=1}^K \begin{pmatrix} -R_{2,i}^S & -\omega_i^S & 0 & -\pi J \\ \omega_i^S & -R_{2,i}^S & \pi J & 0 \\ 0 & -\pi J & -R_{2,i}^S & -\omega_i^S \\ \pi J & 0 & \omega_i^S & -R_{2,i}^S \end{pmatrix} + K \otimes I_4$ | $\bigoplus_{i=1}^K \begin{pmatrix} -R_{2,i}^{MQ} & \omega_i^I - \omega_i^S & 0 & 0 \\ -\omega_i^I + \omega_i^S & -R_{2,i}^{MQ} & 0 & 0 \\ 0 & 0 & -R_{2,i}^{MQ} & \omega_i^I + \omega_i^S \\ 0 & 0 & -\omega_i^I - \omega_i^S & -R_{2,i}^{MQ} \end{pmatrix} + K \otimes I_4$ | $\bigoplus_{i=1}^K \begin{pmatrix} -R_{2,i}^S & -\omega_i^S & 0 & -\pi J \\ \omega_i^S & -R_{2,i}^S & \pi J & 0 \\ 0 & -\pi J & -R_{2,i}^S & -\omega_i^S \\ \pi J & 0 & \omega_i^S & -R_{2,i}^S \end{pmatrix} + K \otimes I_4$ |
| $P_{180,A}^x$ | $I_k \otimes \text{diag}(1, -1, -1, 1)$ | | $I_k \otimes \text{diag}(1, -1, -1, 1)$ |
| $P_{180,A}^y$ | | | $I_k \otimes \text{diag}(-1, 1, 1, -1)$ |
| $P_{180,B}^x$ | $I_k \otimes \text{diag}(1, 1, -1, -1)$ | $I_k \otimes \begin{pmatrix} 0 & 0 & 1 & 0 \\ 0 & 0 & 0 & -1 \\ 1 & 0 & 0 & 0 \\ 0 & -1 & 0 & 0 \end{pmatrix}$ | $I_k \otimes \text{diag}(1, 1, -1, -1)$ |
| $G_{INEPT}$ | $e^{L_{free,A}T} P_{180,A}^x e^{L_{free,A}T}$<br>($T = 1/4J$) | $e^{L_{free,A}T}$<br>($T = 1/2J$) | $\left[ (G_x G_y)^2 (G_y G_x)^2 \right]^2$<br>($G_x = e^{L_{free,A}T} P_{180,A}^x e^{L_{free,A}T}$, $G_y = e^{L_{free,A}T} P_{180,A}^y e^{L_{free,A}T}$,<br>$T = 1/32J$) |
| $R_{AB}$ | $I_k \otimes \text{diag}(0, 0, 1, 0)$ | $I_k \otimes \begin{pmatrix} 0 & 0 & 0 & -1/2 \\ 0 & 0 & 1/2 & 0 \\ 0 & 0 & 0 & 1/2 \\ 0 & 0 & -1/2 & 0 \end{pmatrix}$ | $I_k \otimes \text{diag}(0, 0, 1, 0)$ |
| $R_{BA}$ | $I_k \otimes \begin{pmatrix} 0 & 0 & 0 & 0 \\ 0 & 0 & 0 & 0 \\ 0 & 0 & 0 & 0 \\ 0 & 0 & 1 & -i \end{pmatrix}$ | $I_k \otimes \begin{pmatrix} 0 & 0 & 0 & 0 \\ 0 & 0 & 0 & 0 \\ -i & -1 & -i & 1 \\ 1 & -i & -1 & -i \end{pmatrix}$ | $I_k \otimes \begin{pmatrix} 0 & 0 & 0 & 0 \\ 0 & 0 & 0 & 0 \\ 0 & 0 & 0 & 0 \\ 0 & 0 & 1 & -i \end{pmatrix}$ |
| $M_+$ | $I_k \otimes (1 \ 0 \ 0 \ 0)$ | $I_k \otimes (0 \ 1 \ 0 \ 0)$ | $I_k \otimes (1 \ 0 \ 0 \ 0)$ |
| $M_{obs}$ | $M_+ G_{INEPT} R_{BA} e^{\frac{L_{free,B}t_1}{2}} P_{180,B} e^{\frac{L_{free,B}t_1}{2}} R_{AB} G_{INEPT} \rho_0$ | $M_+ G_{INEPT} R_{BA} e^{\frac{L_{free,B}t_1}{2}} P_{180,B} e^{\frac{L_{free,B}t_1}{2}} R_{AB} G_{INEPT} \rho_0$ | $M_+ G_{INEPT} R_{BA} e^{\frac{L_{free,B}t_1}{2}} P_{180,B} e^{\frac{L_{free,B}t_1}{2}} R_{AB} G_{INEPT} \rho_0$ |

**Figure S1.**
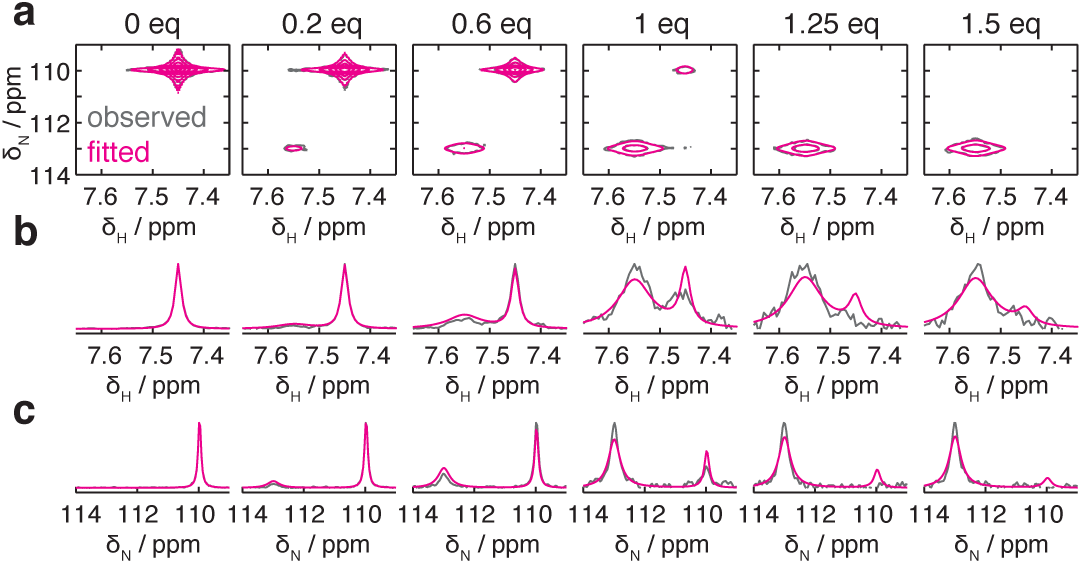
Comparison of one-dimensional and two-dimensional lineshape fitting showing effect of differential relaxation. (**a**) ^1^H,^15^N-HSQC titration data was simulated, including noise, for a two-state binding interaction (*K*_d_ 1 μM, *k*_off_ 5 s^−1^) between two states with different relaxation rates (grey). The results of two-dimensional lineshape fitting (magenta) were in excellent agreement with the true parameters (best-fit *K*_d_ 1.02 μM, *k*_off_ 5.06 s^−1^) (**b,c**) In contrast, simultaneous one-dimensional fits (magenta) to ^1^H and ^15^N cross-sections (grey) did not converge to the observed lineshapes (best-fit *K*_d_ 6.8 μM, *k*_off_ 5.5 s^−1^).

**Figure S2.**
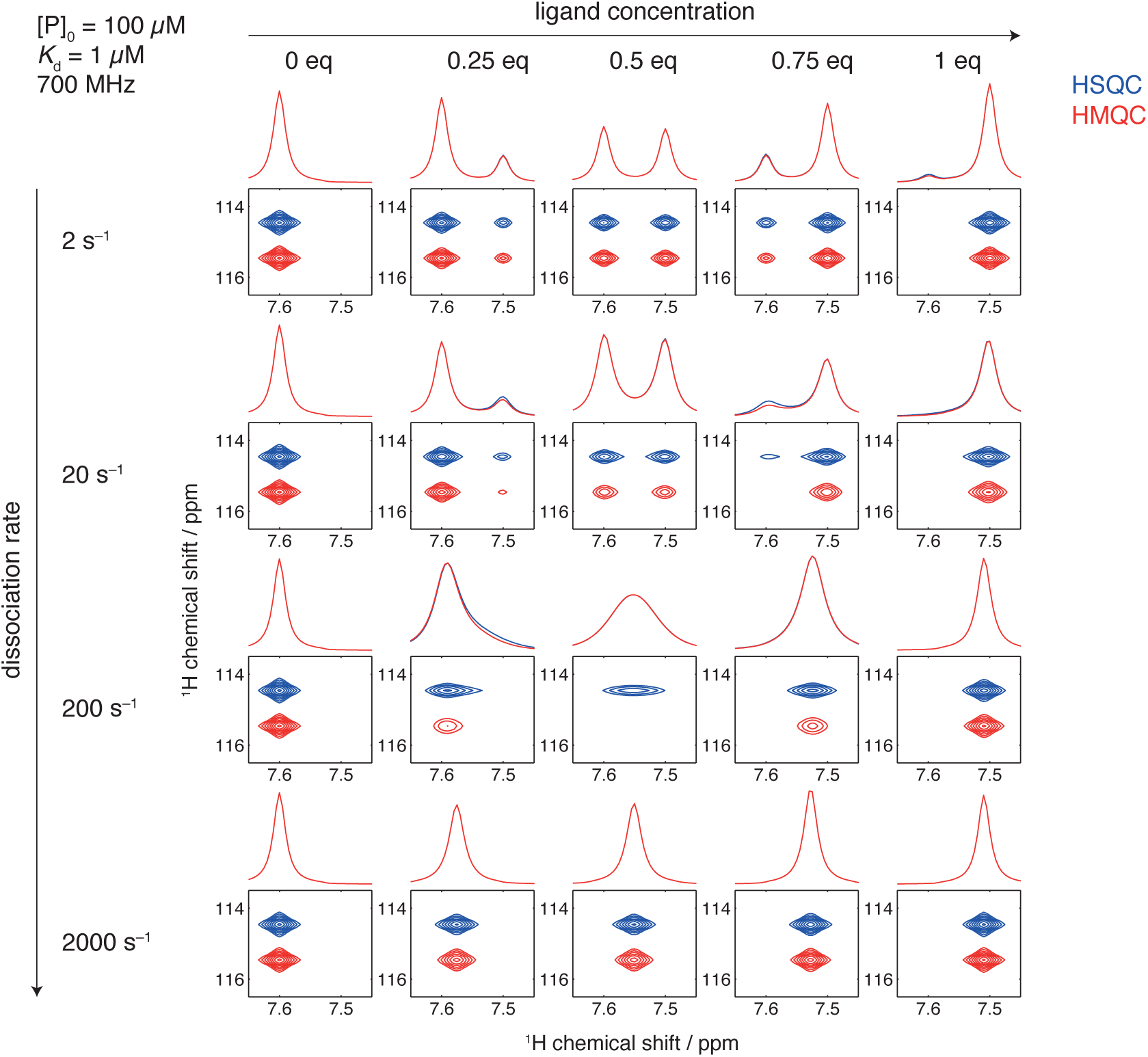
Comparison of simulated exchange lineshapes in ^1^H,^15^N-HSQC and ^1^H,^15^NHMQC experiments for a titration of a protein with an interacting ligand (*K*_d_ = 1 μM, protein concentration [P]_0_ = 100 μM, ^1^H Larmor frequency = 700 MHz, and dissociation rates and ligand concentrations as indicated), illustrating the increased broadening in HMQC spectra due to exchange of multiple-quantum coherences during the indirect evolution period. Normalised cross-sections are also shown, highlighting the effect of differential broadening on lineshapes under conditions of asymmetric slow–intermediate exchange.

**Figure S3.**
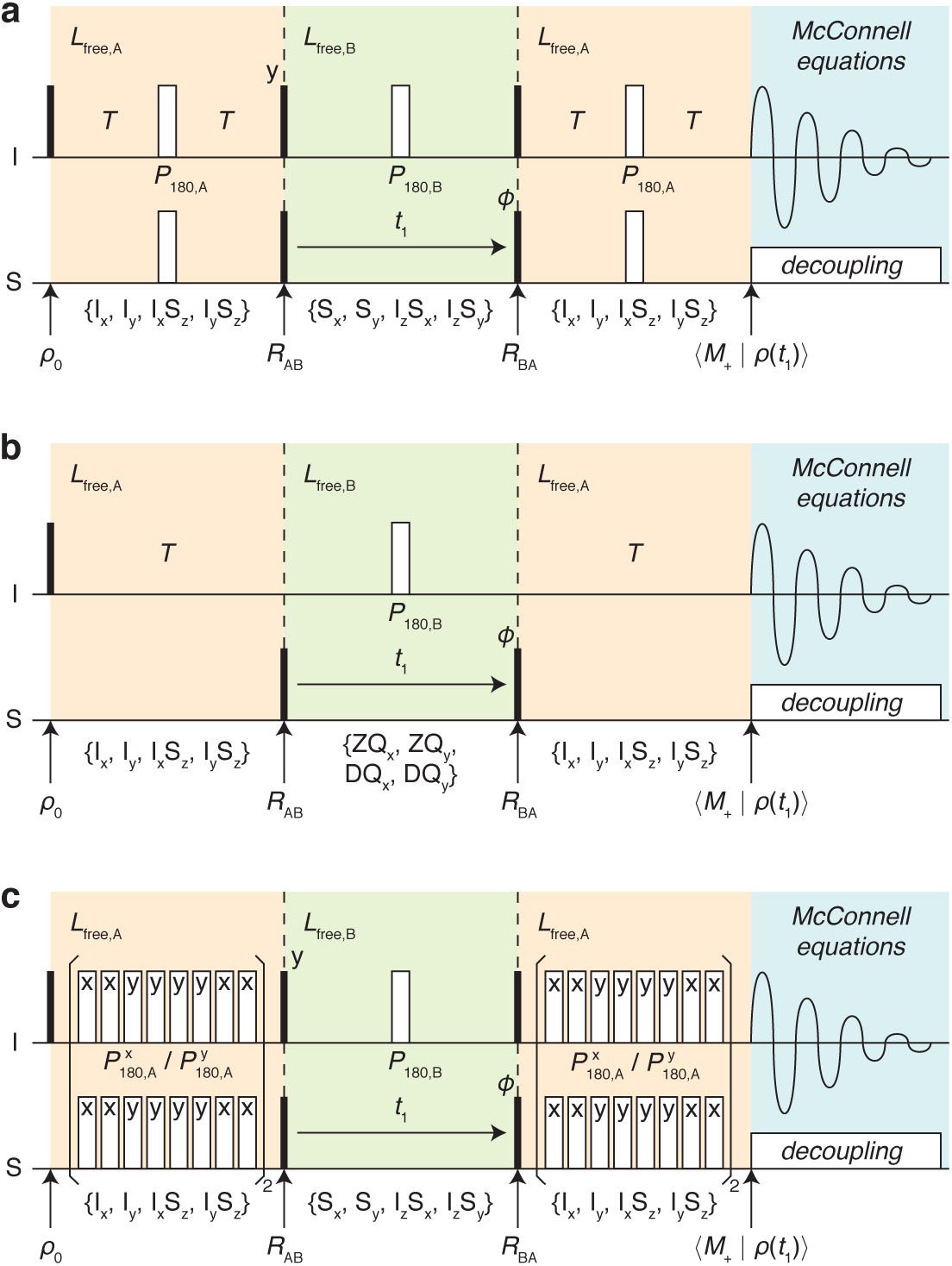
Schematic illustration of calculation schemes detailed in Table S1 for the efficient simulation of two-dimensional spectra obtaining using (**a**) HSQC, (**b**) HMQC, and (**c**) CPMGHSQC pulse sequences. Solid bars indicate 90° pulses and hollow bars 180° pulses, with phase ‘x’ unless otherwise indicated. The initial density operator *ρ*_0_ is propagated through the sequence in the reduced basis spaces indicated with braces and coloured shading, with the superoperators *R* indicating rotations between basis subspaces. The populations of in-phase proton magnetization are determined by projection (onto *M*_+_) at the point of acquisition, with efficient calculation of lineshapes in the direct dimension subsequently obtained through solution of the classical Bloch-McConnell equations.

**Figure S4.**
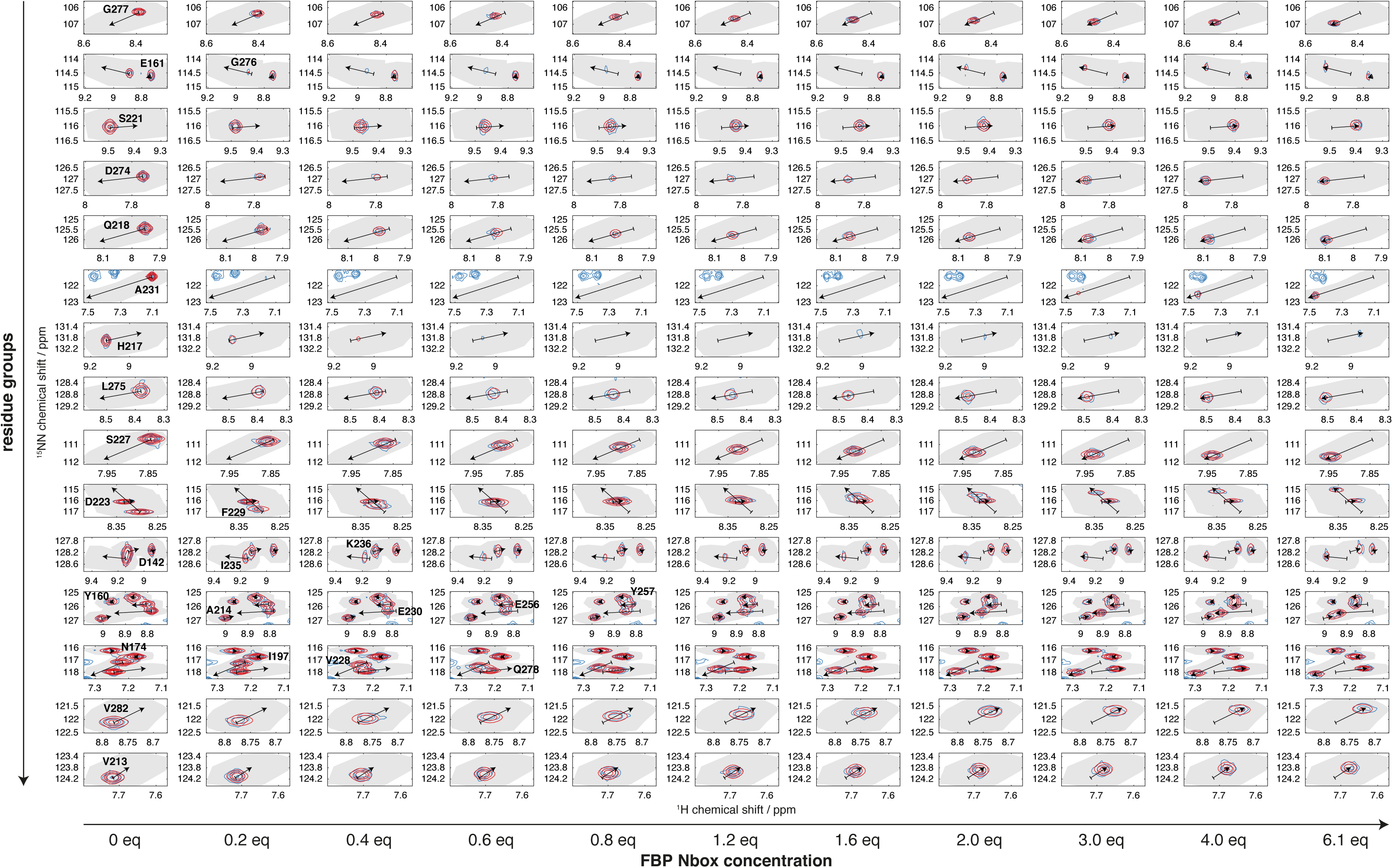
Results of two-dimensional lineshape analysis for the titration of FIR RRM1-RRM2 with FBP Nbox. Blue, observed; red, fitted. Shaded areas indicate the selected regions of interest (ROIs). The chemical shifts of free and bound states determined by the fitting procedure are marked by the tail and head of the arrows.

**Figure S5.**
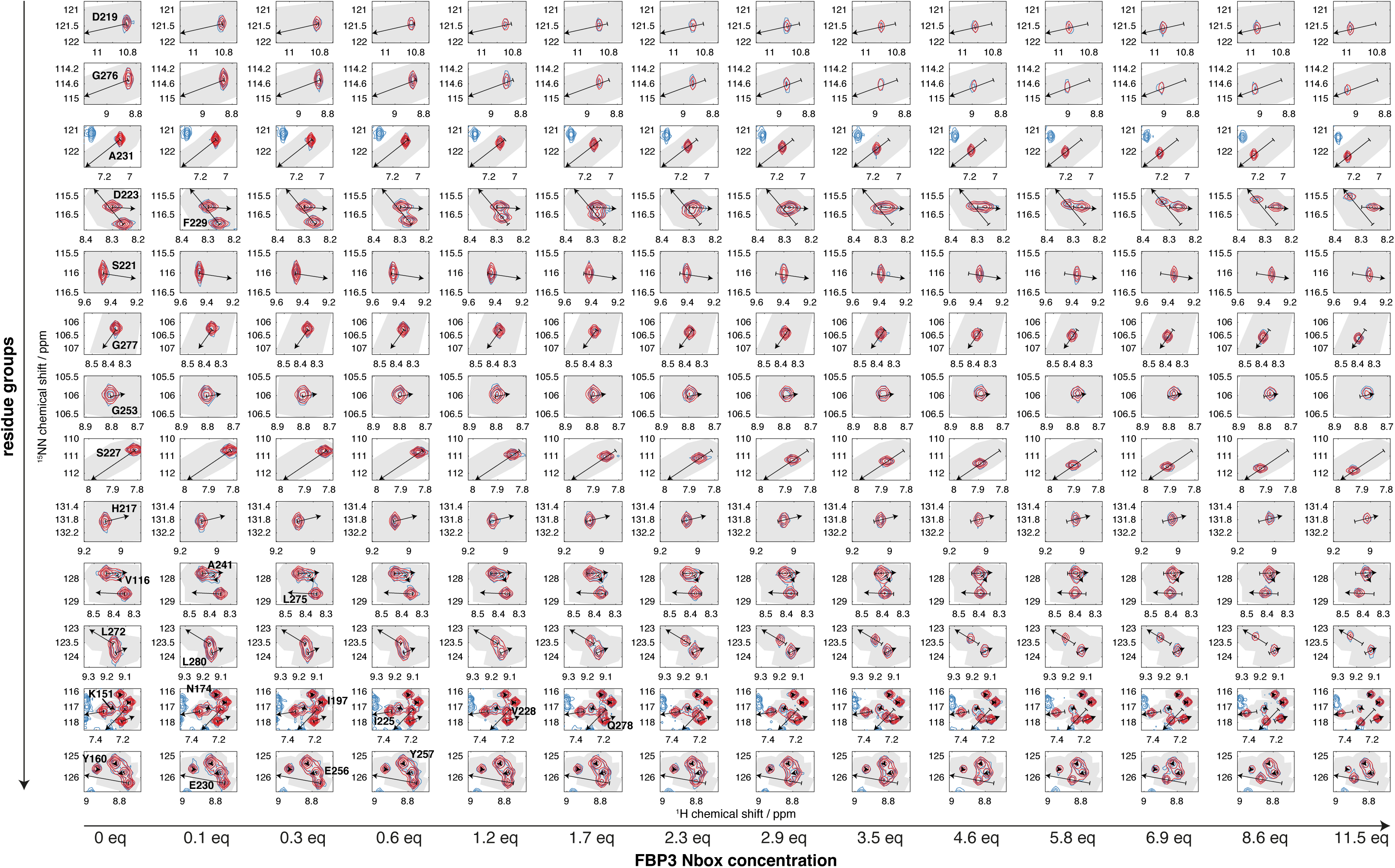
Results of two-dimensional lineshape analysis for the titration of FIR RRM1-RRM2 with FBP3 Nbox. Blue, observed; red, fitted. Shaded areas indicate the selected regions of interest (ROIs). The chemical shifts of free and bound states determined by the fitting procedure are marked by the tail and head of the arrows.

**Figure S6.**
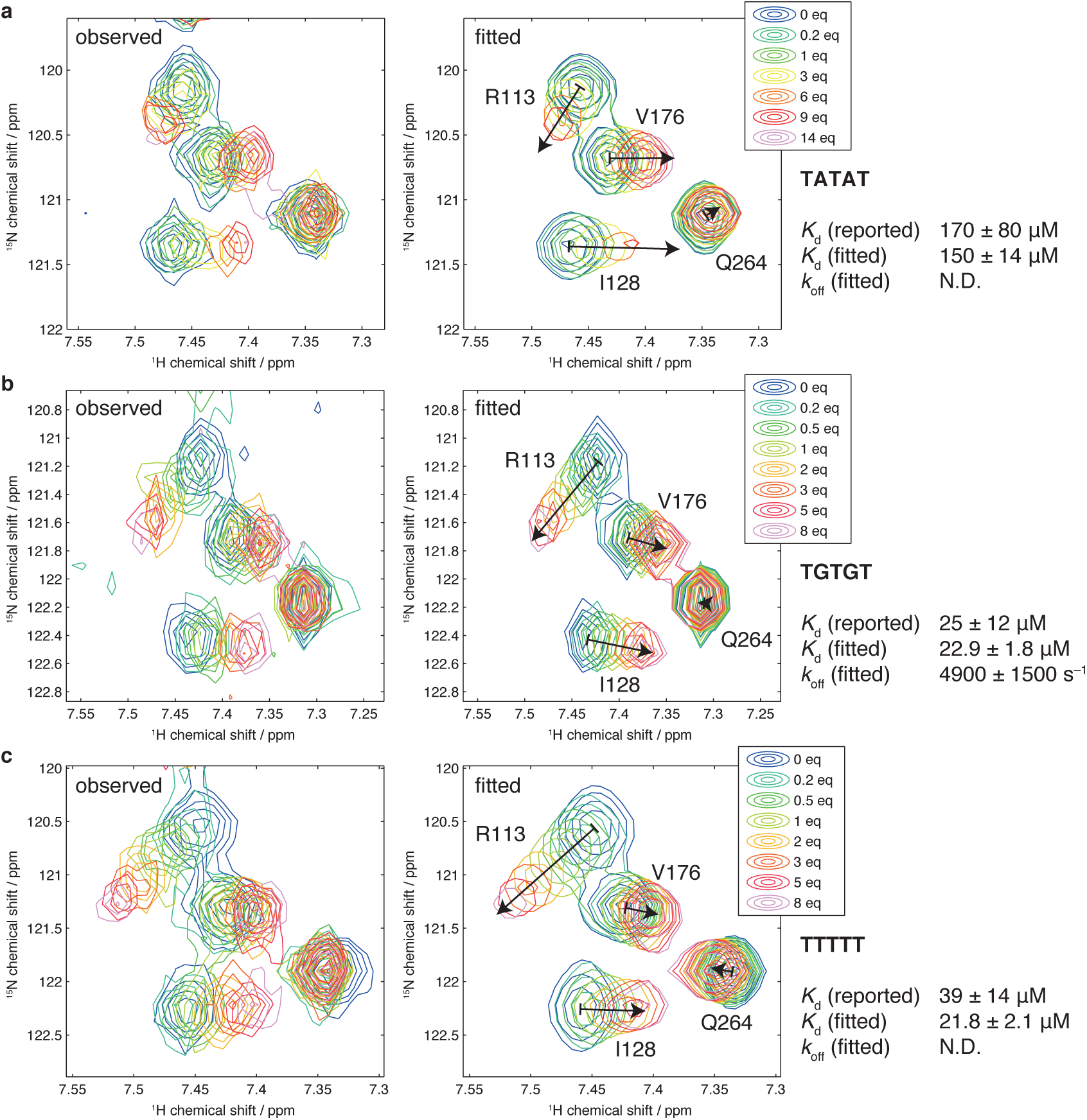
Results of two-dimensional lineshape analysis for titrations of FIR RRM1-RRM2 with the oligonucleotides (**a**) TATAT, (**b**) TGTGT, and (**c**) TTTTT. The chemical shifts of free and bound states determined by the fitting procedure are marked by the tail and head of the arrows shown in the fitted spectra.

**Figure S7.**
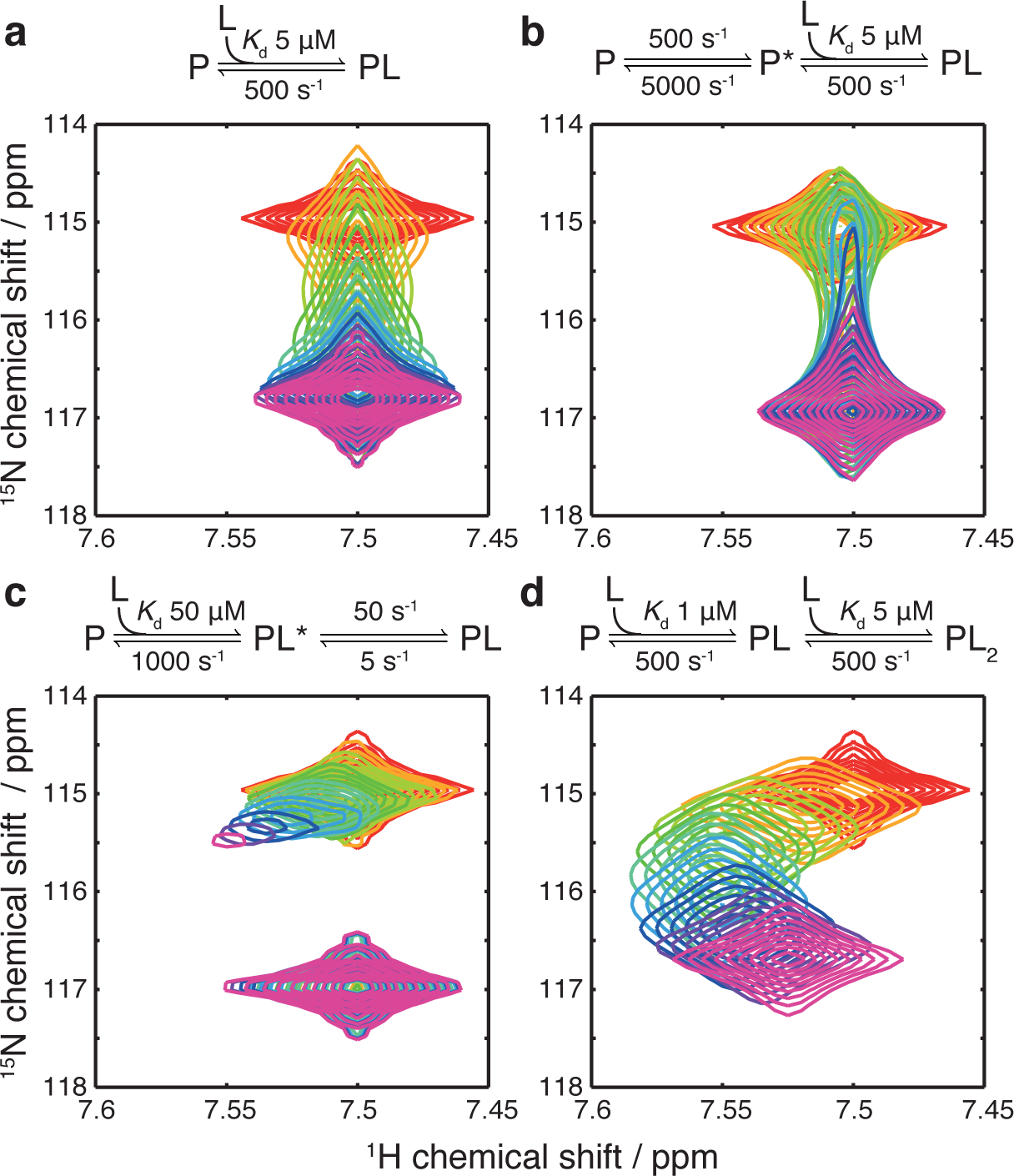
Simulation of a variety of binding mechanisms to illustrate the potential for 2D NMR measurements to discriminate between different mechanistic scenarios: (**a**) simple two-state association; (**b**) conformational selection; (**c**) induced fit; and (**d**) two sequential association reactions. ^1^H,^15^N-HSQC spectra are shown simulated at 700 MHz, 50 μM protein concentration, 0 μM (red) to 100 μM (purple) ligand concentrations. Dissociation constants and rate constants are indicated on the reaction schemes above each panel.

**Figure S8.**
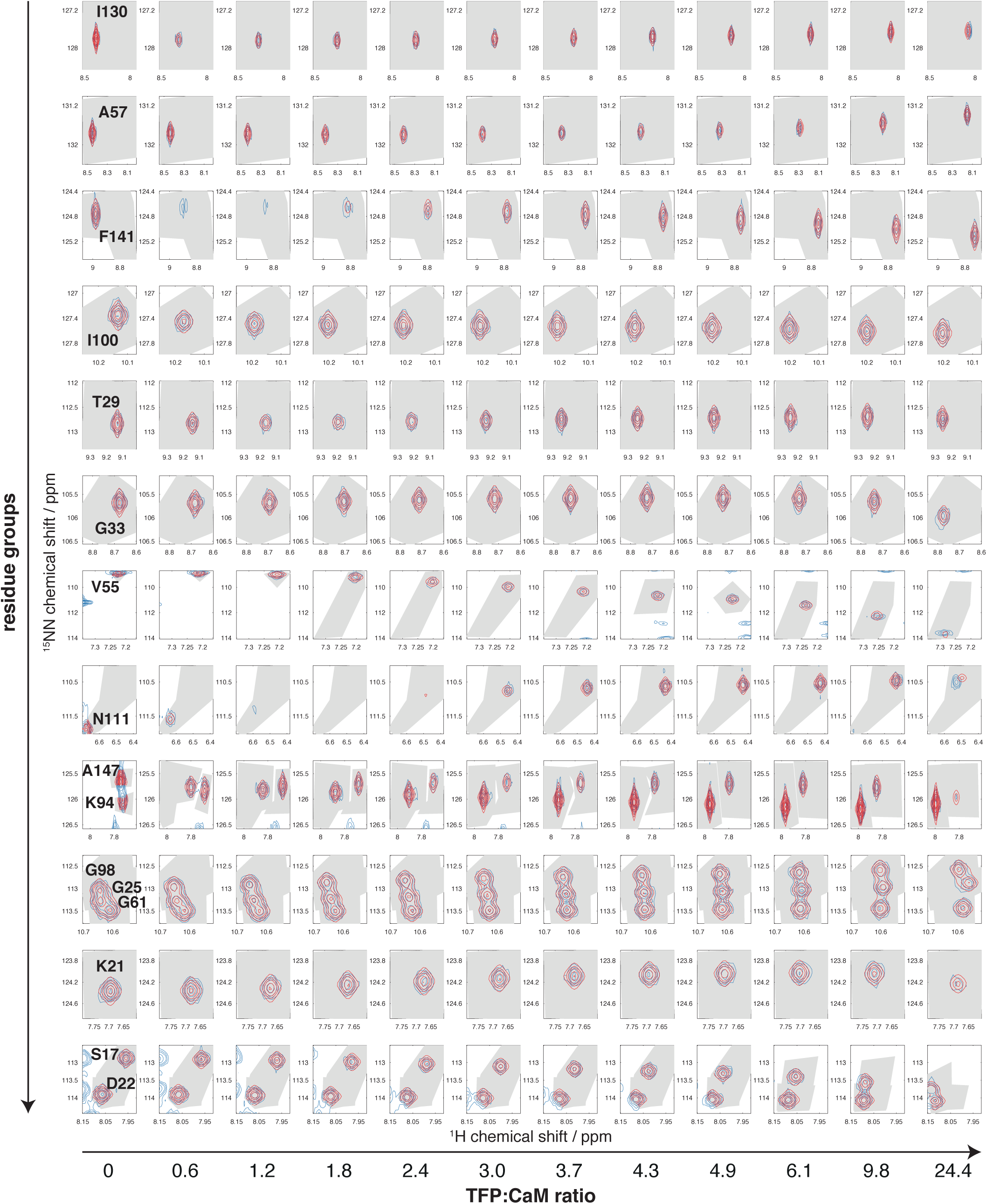
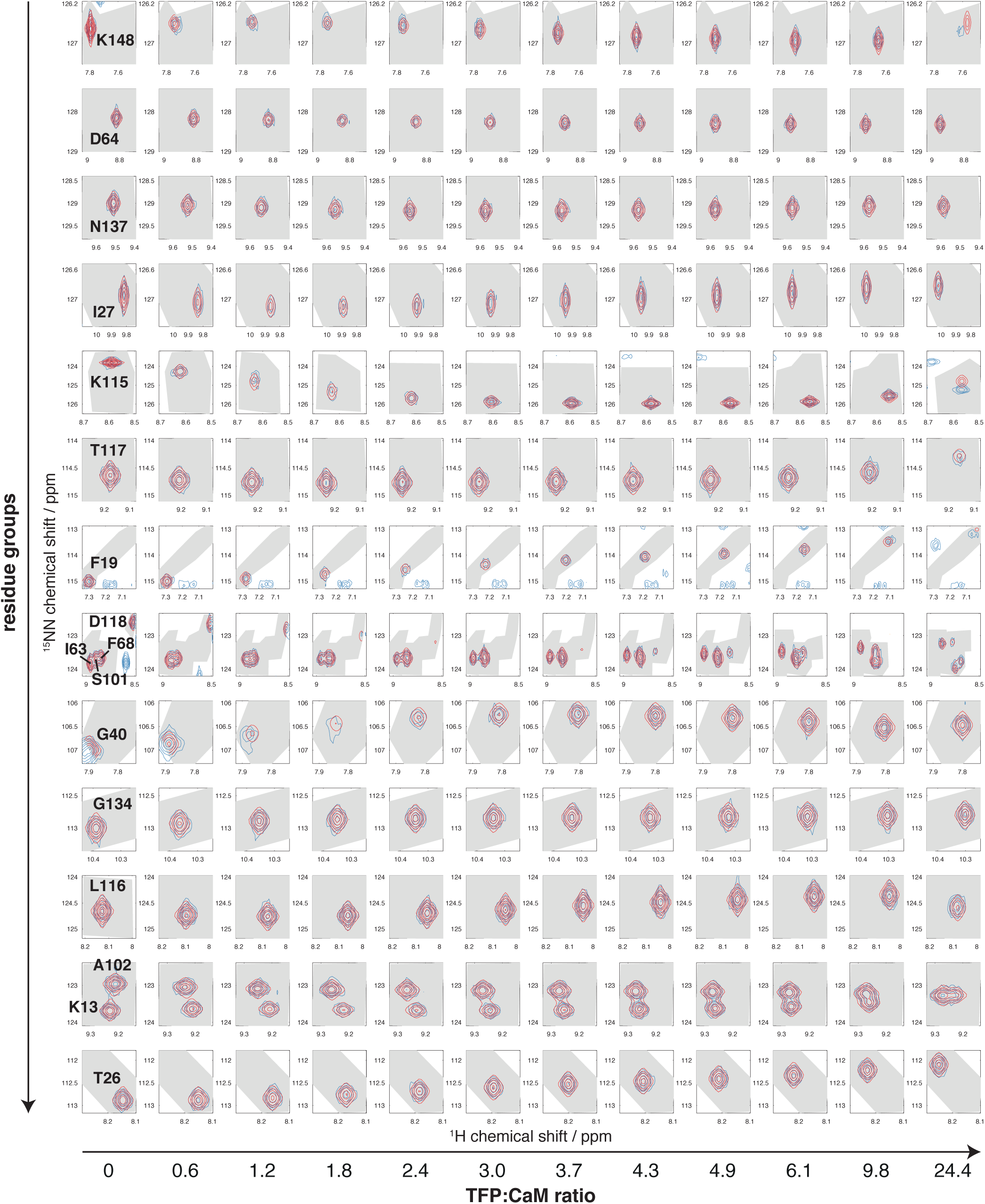
Results of two-dimensional lineshape analysis for the titration of Ca^2+^-CaM with TFP. Blue, observed; red, fitted. Shaded areas indicate the selected regions of interest (ROIs).

**Figure S9.**
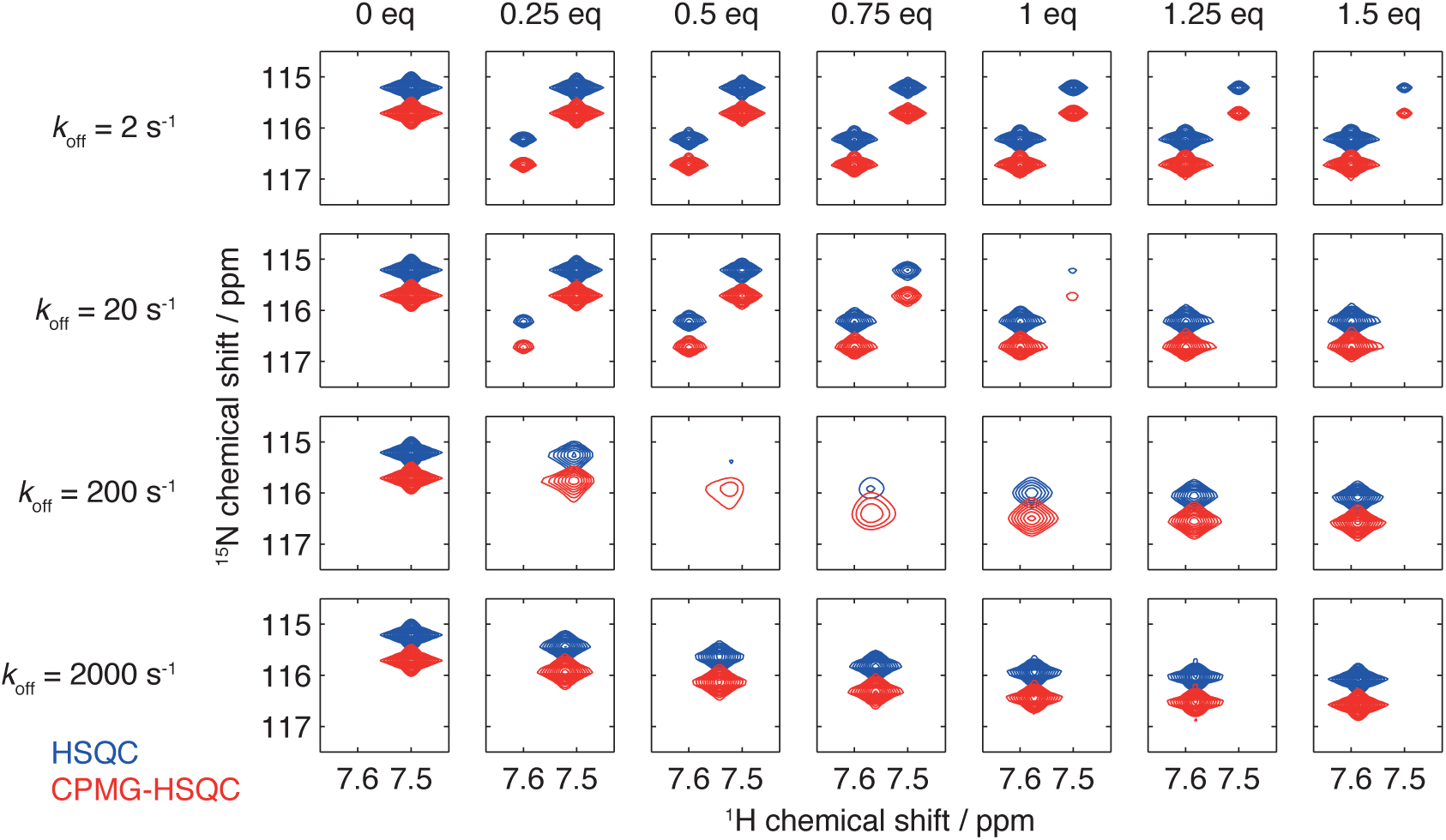
Comparison of HSQC (blue) and CPMG-HSQC (red) pulse sequences for a simulated two-state binding reaction (50 μM protein concentration, 5 μM *K*_d_, 700 MHz ^1^H Larmor frequency) with dissociation rates and ligand concentrations as indicated. Contour levels are fixed across all spectra.

